# Effects of prefrontal theta burst stimulation on neuronal activity and subsequent eating behavior: an interleaved rTMS and fNIRS study

**DOI:** 10.1101/2020.09.08.287896

**Authors:** Idris Fatakdawala, Hasan Ayaz, Adrian Safati, Mohammad Nazmus Sakib, Peter A. Hall

## Abstract

The dorsolateral prefrontal cortex (dlPFC) and dorsomedial prefrontal cortex (dmPFC) are important nodes for self-control and decision-making, but through separable processes (cognitive control versus evaluative processing). This study aimed to examine the effects of excitatory brain stimulation (intermittent theta-burst stimulation; iTBS) targeting the dlPFC and dmPFC on food choice. iTBS was hypothesized to decrease consumption of appetitive snack foods, via enhanced interference control for dlPFC stimulation and reduced delay discounting for dmPFC stimulation. Using a single-blinded, between-subjects design, participants (*N* = 43) were randomly assigned to of the three conditions: 1) iTBS targeting the left dlPFC, 2) iTBS targeting bilateral dmPFC, or 3) sham. Participants then completed two cognitive tasks (delay discounting (DD) and Flanker), followed by a taste test. fNIRS imaging revealed increases in medial PFC activity were evident in the dmPFC stimulation group during the DD task; likewise, a neural efficiency effect was observed in the dlPFC stimulation group during the Flanker. Gender significantly moderated consumption during the taste test, with females in the dmPFC showing paradoxical increases in food consumption compared to sham. Findings are consistent with possible amplification of positive evaluative processing in the presence of dietary restraint, vis-à-vis excitation of the mPFC.

## Introduction

The prefrontal cortex (PFC) contains several important nodes of the executive control network, supporting a variety of functions including inhibitory control and evaluative processing (MacPherson *et al*., 2002; Siddiqui *et al*., 2008). Dietary self-control is partially dependent on the lateral PFC, as it is thought to facilitate suspension of prepotent approach behaviors to high caloric foods in the interests of restraint goals (Hall, 2016; Lowe *et al*., 2019). For this reason, it has been hypothesized that the extent to which indulgent eating occurs in permissive contexts may be partially dependent on the integrity of the lateral prefrontal cortex as well as its functional connectivity with other brain regions involved in value and salience processing (Hall, 2016; Donofry *et al*., 2019; Lowe *et al*., 2019). There is an accumulating body of literature linking attenuated dlPFC function with indulgent eating (Brooks *et al*., 2013; Fuguo Chen *et al*., 2018; Han *et al*., 2018; Schmidt *et al*., 2018; Donofry *et al*., 2019). Likewise, some evidence indicates that indulgent food consumption is linked with attenuated function of these same cortical nodes (Kalon *et al*., 2016; Rui Chen *et al*., 2018), supporting the possibility of a reciprocal relationship between PFC function and obesity (Lowe *et al*., 2019). A meta-analysis of experimental studies examining the effects of non-invasive brain stimulation methods revealed that excitatory stimulation of the dlPFC decreases, and suppressive stimulation increases, indulgent food consumption (Lowe *et al*., 2017).

In the clinical trial literature, multi-session rTMS intervention studies have shown that excitatory stimulation of the left dlPFC reduces cravings for appetitive foods among those with eating disorders (Uher *et al*., 2005; Van den Eynde *et al*., 2010). Likewise, multi-session excitatory rTMS targeting the left dlPFC was found to be effective in enhancing weight loss and decreasing food intake in obese patients in at least one recent study (Kim *et al*., 2019). On the other hand, in the experimental literature, continuous theta burst stimulation (cTBS; a suppressive variant of rTMS) targeting the left dlPFC results in amplified cravings and consumption of similar foods when applied in single-session format (Lowe *et al*., 2014; Lowe, Staines, *et al*., 2018). Meta-analyses examining the effects of non-invasive brain stimulation techniques reveal medium effect sizes on food cravings (Jansen *et al*., 2013; Lowe *et al*., 2017) and consumption outcomes (Hall *et al*., 2017; Song *et al*., 2019), both in favour of active over sham stimulation. Such findings are similar to those observed for neuromodulation studies involving other types of cravings (Jansen *et al*., 2013).

Given that brain stimulation protocols targeting the dlPFC reliably alter task performance in theorized directions (Lowe, Manocchio, *et al*., 2018), it is reasonable to posit that excitatory stimulation effects on eating might be mediated through improved performance on cognitive interference tasks that load heavily on inhibitory functions. However, the specific PFC subregion targeted is an important consideration. Like executive control, evaluative processing—supported by the dmPFC—is also implicated in both indulgent eating (Hare *et al*., 2011; Rangel, 2013; Dietrich *et al*., 2016; Berkman *et al*., 2017; Rui Chen *et al*., 2018; Hall, 2019) and self-regulatory processes in relation to tempting stimuli (Cho *et al*., 2015; Hall, 2019). Functional connectivity between the dmPFC and other brain regions is altered following gastric bypass surgery, one of the more successful approaches to radical weigh reduction (Li *et al*., 2018)

For these reasons, stimulation of the dmPFC might also impact indulgent eating by altering evaluative processing of food-related stimuli or affecting the link between value processing and eating behavior regulation. The directionality of anticipated effects on eating indulgence are clear for the lateral PFC, given its role in executive control; however, there is less certainty about the dmPFC, given that it is implicated in value computation and mind wandering, both of which could sway behavior both toward or away from eating indulgence, depending on the preponderance of active value signals or sensory stimuli (in favor of indulgent consumption, or away from it). In either case, the medial PFC, broadly speaking, appears to be implicated in value signal processing in a manner that is broadly relevant for self-regulation of eating behavior (Hare *et al*., 2011; Rangel, 2013; Berkman *et al*., 2017).

In this context, the purpose of the current study was to examine the role of two PFC sub-regions—the dlPFC and dmPFC—in eating behavior, as mediated through either inhibitory or evaluative processing. fNIRS was used to examine task-related neural activity in both PFC subregions following rTMS (Curtin, Tong, *et al*., 2019), and eating indulgence was assessed using a bogus taste test paradigm involving calorie-dense snack foods. Given research precedent, we anticipated that lateral PFC excitatory stimulation would enhance neural activity within the dlPFC during Flanker performance, and subsequently reduce indulgent eating during the bogus taste test. Hypotheses regarding dmPFC stimulation were more tentative, given the less certain effect of excitatory TBS protocols on neural activity in this PFC subregion, and the potentially agnostic role of evaluative processing in the self-regulatory process in the eating context. Finally, given that gender differences exists in food cravings (Hallam *et al*., 2016), along with possible differences in brain stimulation effects between younger and older adults (Oberman and Pascual-Leone, 2013; Gutchess, 2014), we also examined for participant gender and age group as moderators for our experimental outcomes.

## Methods

### Participants

A sample of 43 participants were recruited for the study. To assess the differences in the effects of iTBS by age, the recruitment was stratified as follows: 22 younger adults and 21 middle-to-older aged adults (“older adults”). Younger adults were between 18-30 years of age and were all recruited from advertisements placed around campus. The older adults were between 40-75 years of age and were recruited from campus, an aging participant pool as well as from local community centres. All eligible participants were right-handed, neurologically healthy and naïve to TMS; prior to participation individuals were screened for any physical and neurological conditions that would contraindicate rTMS, using a standard screening form (Rossi *et al*., 2009). Participants with food intolerances or health conditions (e.g. diabetes) that would preclude normative participation in the taste test were also excluded. Following explanation of risks and benefits of participation, electronic informed consent was obtained from all participants prior to the start of the study. In exchange for their participation, each participant received a $25 gift card as reimbursement for their time. The study was reviewed by and received ethics clearance from the institutional research ethics committee.

### Procedure

The study employed a single-blinded between-subject design in which participants were randomly assigned to one of the three conditions: iTBS targeting the dlPFC, iTBS targeting the dmPFC and sham iTBS targeting the vertex. Participants were asked to refrain from eating or consuming any caffeinated beverages 3 hours prior to the study; adherence to these requirements was checked with the completion of the consent and screening forms. All computer tasks were presented using the Inquisit (Millisecond Software) on a 27-inch monitor. Prior to commencing the computer tasks, participants were asked to follow the instructions that were presented on the monitor and respond as quickly and accurately as possible while completing each cognitive task. The ambient lighting and temperature conditions were stable across all participants.

Following consent procedures, resting motor threshold was determined followed by active or sham stimulation, as randomly assigned. Ten minutes post-stimulation, participants were asked to complete the two computerized cognitive tasks in the following order: three blocks of the delay discounting task, followed by four blocks of the flanker inhibition task. Between each block, there was a 15 second rest period. While performing these tasks, changes in blood oxygenation levels were measured using the fNIRS protocol (see below). Following the completion of these tasks, approximately 30 minutes post stimulation (when iTBS excitatory effects would be maximal), the participants were given the opportunity to sample five different calorie dense snack foods under the guise of examining the relationship between brain function and taste perception. Change in the weight of the food from pre-to-post tasting was measured surreptitiously to quantify the amount of food that was consumed.

### Brain Stimulation Protocol

iTBS targeting the left dlPFC was administered using a 75mm figure-8 coil (MCF-B65), while the iTBS stimulation for the bilateral dmPFC was administered using a 97cm figure-8 coil (MCF-B70), both which were connected to a Mag Pro x100 biphasic stimulation unit. Sham stimulation was delivered with a placebo version of the MCF-B65 (i.e., MCF-P-B65), targeting the vertex. The iTBS stimulation intensity to the left dlPFC was individually calibrated based on each participant’s resting motor threshold (RMT), as per standard practise. The RMT was defined as the lowest stimulation intensity required to induce a motor evoked potential (MEP) in the right abductor pollicus brevis muscle (the right thumb muscle) of > 50 uV peak-to-peak amplitude, respectively, in 5/10 consecutive trials of stimulating the motor cortex. In order to guide coil placement for stimulating both the motor cortex (for determining the RMT) and the cortical target regions of interest, an EEG cap with electrodes arranged in the international 10-20 system fitted according to standard anatomical landmarks. The determination of the RMT was made using the C3 position to approximate the motor cortex; the iTBS stimulation site for the left dlPFC was approximated by the F3 electrode position. Finally, the iTBS stimulation site for the bilateral dmPFC was approximated as 2/3 of the distance from the nasion to the vertex, as per prior research precedent (Berkers *et al*., 2017).

Following RMT determination, the intensity for left dlPFC stimulation was set at 80% of the RMT. Stimulation consisted of triplet stimuli applied in the theta burst pattern (three 50 Hz pulses repeated at a frequency of 5 Hz); TBS was applied for 2 seconds for every 10 second period (i.e. 2 seconds of TBS followed by 8 seconds of rest), for a duration of 190 seconds, totaling 600 stimuli (Huang *et al*., 2005). Participants assigned to the sham iTBS condition received a procedure identical to that described above using the placebo version of the same coil (MCP-P-B65) targeting the vertex (Cz position) instead. The MCF-P-B65 blocks 80% of the stimulation intensity delivered by the coil, which is otherwise identical to the MCF-B65. In the dmPFC stimulation condition, stimulation intensity was set at a fixed intensity of 30% of the maximal stimulator output in keeping with prior research using this stimulation site (Berkers *et al*., 2017).

### Functional Near Infrared Spectroscopy Protocol

Functional Near Infrared Spectroscopy (fNIRS) is a non-invasive, optical neuroimaging technique that uses near-infrared (NIR) light sources and detectors to quantify changes in blood oxygenation levels within cortical brain tissues following neuronal activation (Ferrari and Quaresima, 2012; Curtin and Ayaz, 2018; Pinti *et al*., 2020). In order to measure cerebral activation, fNIRS relies on a hemodynamic response, which produces a relative increase in oxygenated hemoglobin (HbO) and decrease deoxygenated hemoglobin (HbR) during neuronal activity (Ayaz *et al*., 2019; Pinti *et al*., 2020). Because HbO and HbR absorb NIR light at different wavelengths, fNIRS is able to take advantage of chromophoric features of hemoglobin to detect changes in brain activation.

For this study, fNIR Devices 203C imaging system was used to quantify regional oxygen saturation in the prefrontal cortex. The equipment device consisted of a headband with a sensor pad, which was embedded with 4 LED light sources and 10 light detectors, joined to create 16 channels (plus two short channels). Two wavelengths of light, at 730nm (for HbR) and 850nm (for HbO) were measured using the COBI Studio software (Ayaz *et al*., 2011). Signal processing procedure following (Ayaz *et al*., 2012), briefly: for each participant, raw fNIRS light intensity data were low-pass filtered with a finite impulse response, linear phase filter with order 20 and cut-off frequency of 0.1 Hz to attenuate the high frequency noise, respiration and cardiac cycle effects. Each participant’s data was checked for any potential saturation (when light intensity at the detector was higher than the analog-to-digital converter limit) and motion artifact contamination by means of a coefficient of variation-based assessment (Ayaz *et al*., 2010). fNIRS data for each task block were extracted using beginning and end timings based on the experiment and hemodynamic changes for each 16 brain areas during each trial block were calculated separately using the Modified Beer Lambert Law (MBLL) with respected to the local baseline at the beginning of the respective block. The hemodynamic response at each optode was then averaged across time for each trial block to provide a mean hemodynamic response at each optode for each block to be used in statistical analysis.

### Cognitive tasks

#### Delay Discounting Task

Participants were asked to complete a variant of the delay discounting paradigm described by Koffarnus and Bickel (Koffarnus and Bickel, 2014). Participants were presented with three blocks, with the option of choosing between two choices: a fixed larger commodity for which the delay was adjusted from trial to trial verses a fixed smaller commodity that was immediately available. The magnitude of the immediately available option for each block was set at half of the delayed option (Block 1: $5 vs. $10, Block 2: $500 vs. $1000, Block 3: $500,000 vs. $1,000,000). Each block consisted of five trials. The first-choice trial for each block was always set with the larger commodity delayed at 3 weeks. For subsequent trials, the delay for the larger option were adjusted depending on the participant’s previous choice; delay was adjusted up if the delayed choice was chosen or down if the immediate choice was chosen on the previous trial. k values are determined at the end of last trial. A smaller k value is taken to indicate the relative absence of discounting, thereby implying a preference for delayed rewards. A higher *k* value is indicative of a strong discounting rate, thereby implying a preference for immediate rewards.

#### Flanker Task

Participants were asked to complete a modified version of the Eriksen Flanker task, which was used to measure behavioral inhibition. In this task, participants were presented with a stimulus consisting of a set of seven letters and were asked to make directional responses to the letter in the centre (the target stimuli) arranged among an array of flanking letters (nontarget stimuli), by pressing the corresponding keyboard key that is to assigned target stimuli. The target letters “H” and “K” were assigned to either the “A” or “D” keyboard key, while the target letters “S” and “C” were assigned to the alternative key. Participants were presented with two conditions: 1) congruent noise condition, in which the target letter were flanked by the letter corresponding to the same keyboard key response (i.e. HHHKHHH or CCCSCCC). And 2) incongruent noise condition, in which the target letter was flanked by the letters assigned to the other keyboard key response (i.e. CCCHCCC or HHHSHHH). Participants were required to make their response to the target letter as quickly and accurately as possible. The task began with a practice block, (60 trials of incongruent + congruent), followed by 4 blocks (2 blocks of each condition), completed in the following order: 50 trials of the congruent task, 75 trials of the incongruent task, 50 trials of the congruent task and 50 trials of the incongruent task. Flanker interference score was calculated by taking the difference in the latency of the correct trials in congruent noise condition from the incongruent noise condition. A higher score therefore reflects worse performance on the task.

#### Taste Test and Food Ratings

Taste test paradigms are commonly used in the eating literature as a reliable and valid metric of consumption; quantity consumed is positively associated with food palatability (Robinson *et al*., 2017), level of hunger (Robinson *et al*., 2017), and responsive to acute manipulations of executive function using TMS targeting the left dlPFC (Lowe, Staines, *et al*., 2018). In the current version of the paradigm, participants were presented an array of five calorie-dense snack foods (3 types of Pringles potato chips and 2 types of Belgian chocolate balls (milk chocolate and salted chocolate). Participants were given 15 minutes for the task, during which they were asked to complete a 7-item self-reported food ratings questionnaire for each food item presented. Prior to the task, participants given a verbal cue “you can eat as much as you would like while making your taste ratings”, under the guise that the main purpose of the task was to explore TMS effects on sensory processing. The experimental foods were surreptitiously weighted before and after the taste test, where the difference (amount of food consumed) was recorded (in grams) as the primary metric of food consumption. A total score and separate food type (i.e., sweet vs. salty) scores were calculated.

### Statistical Analyses

Descriptive statistics were computed to examine the distribution of each of the continuous outcome variables of interest: i) food consumption, ii) flanker interference scores and iii) averaged delay discounting *k* values by treatment condition. Boxplots were generated to visually examine the characteristics of each variable distribution and identify any outliers. All the outcome variables in the dataset were subject to winsorization to limit the effects of extreme outliers. Four outliers were identified for the food consumption variable: two below the 5^th^ percentile value and two above the 95^th^ percentile value, which were than replaced with an assigned percentile values. In addition, granular analyses was conducted to compare differences in type of food consumed (salty vs. sweet). One participant’s flanker score was dropped because of accuracy scores below chance level. Four additional scores were replaced with winsorized values. The delay discounting variable (*k*) was averaged across the three trials, to obtain an average discounting rate for each participant. Because the variable displayed significant skewness, *k* scores were then subject to a Log10 transformation in order to improve the distributional properties. Four values were replaced with their assigned winsorized percentile values. Next, univariate general linear models were employed to examine the effects of the stimulation condition (active dlPFC stimulation vs. active mPFC stimulation vs. sham stimulation) on the candidate outcome variables (i.e., food consumption, flanker performance, delay discounting). Age category and gender were examined as a moderator of treatment effects. Planned comparisons were conducted using independent t-tests. Finally, fNIRS data from the cognitive tasks were analyzed for differences in oxy hemoglobin concentrations between stimulation conditions for each channel, where heat maps were designed using corresponding *p*-values. Each channel was subject to winsorization prior to running the GLM’s.

## Results

The three stimulation groups did not differ with respect to age (*F*(2,40) = 0.043, *p* = 0.953), gender (*F*(2,40) 0.080, *p* = 0.924), BMI (*F*(2,40) = 1.601, *p* = 0.214) and time of last meal *F*(2,40) = 1.724, *p* = 0.191; Table 1). Likewise, taste ratings did not differ among the three groups with respect to overall appeal (*F*(2,40) = 0.671, *p* = 0.517), saltiness (*F*(2,40) = 0.159, *p* = 0.854), sweetness (*F*(2,40) = 0.651, *p* = 0.546), greasiness (*F*(2,40) = 1.811, *p* = 0.177), healthiness (*F*(2,40) = 0.460, *p* = 0.634) or globally palatability (*F*(2,40) = 0.566, *p* = 0.572); this confirms that stimulation did not influence sensory processing of the foods presented in the taste test.

**Table 1.**
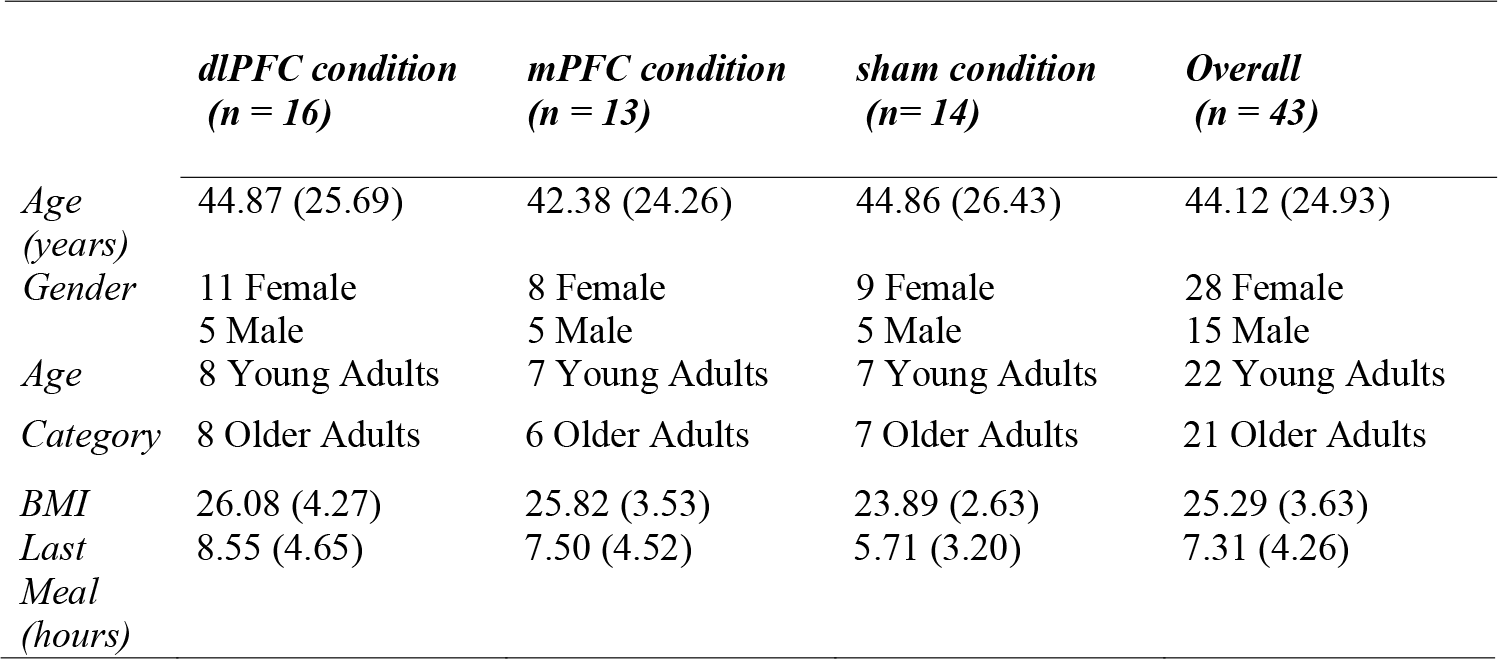
Mean (*SD*) for demographic variables by treatment condition

### Neural Response to iTBS

#### Flanker OxyHb

Channel by channel OxyHb response is displayed using a heat map (Figure 1, left) using corresponding *F* statistic values derived from a contrast between active and sham stimulation groups during the Flanker interference task; the significant contrast is overlaid upon a 3 dimensional greyscale brain (Figure 1, right).

**Figure 1.**
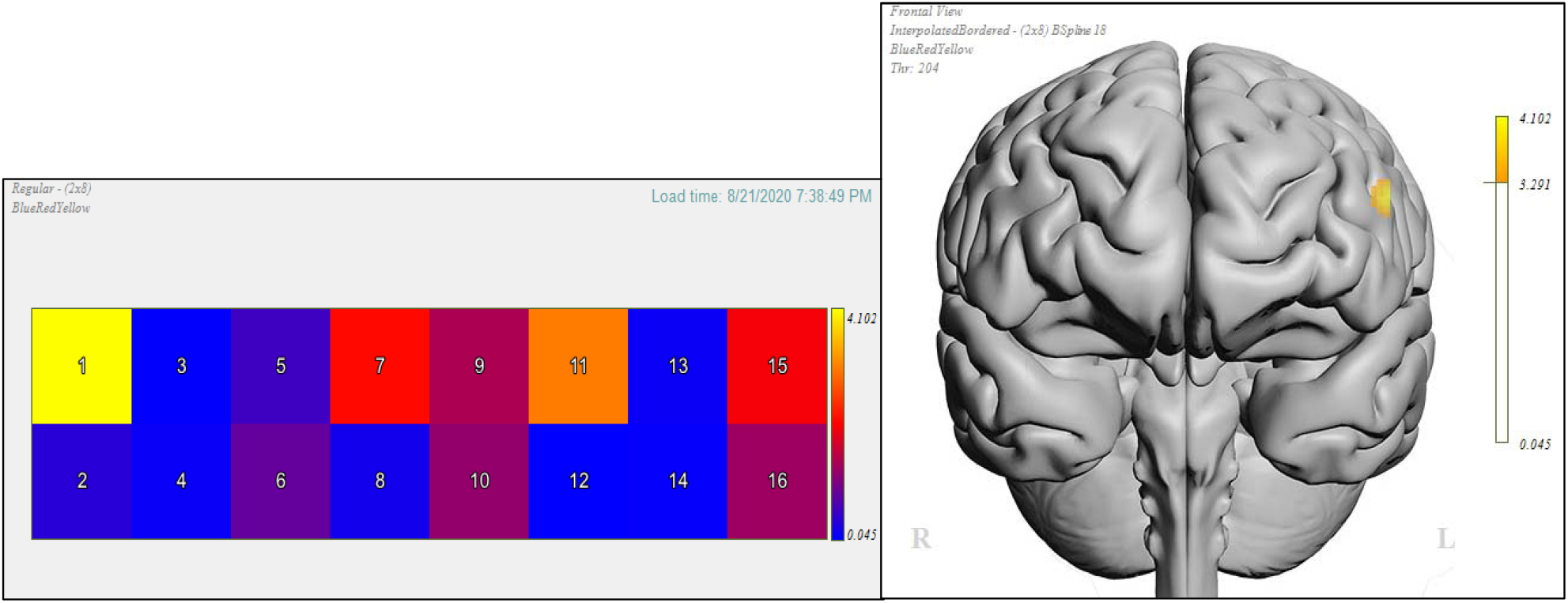
Left: Heat map of channels 1-16 illustrating the difference in oxy-hemoglobin concentration between active iTBS (targeting the left dlPFC) vs. sham iTBS during the Flanker task. Colour coding was represented by the strength of the *p*-values; warmer colours represent stronger active vs. sham contrast; darker shades of blue represent weak to no differences. Right: 3D anatomical overlay of the significant contrast difference between active vs. sham.

A significant incongruent/congruent contrast effect was observed for channel 1 (*F*(2,29) = 4.102, *p* = .027) comparing stimulation conditions (active iTBS vs. sham iTBS). Specifically, those in active iTBS condition had significantly OxyHb response (dlPFC condition = *M* = −.646, *SE* = .386, mPFC condition = M = .146, *SE* = .735) than those in the sham condition (*M* = 2.21, *SE* = 1.02). Variables means for all stimulation conditions have been depicted in Figure 2 below.

**Figure 2.**
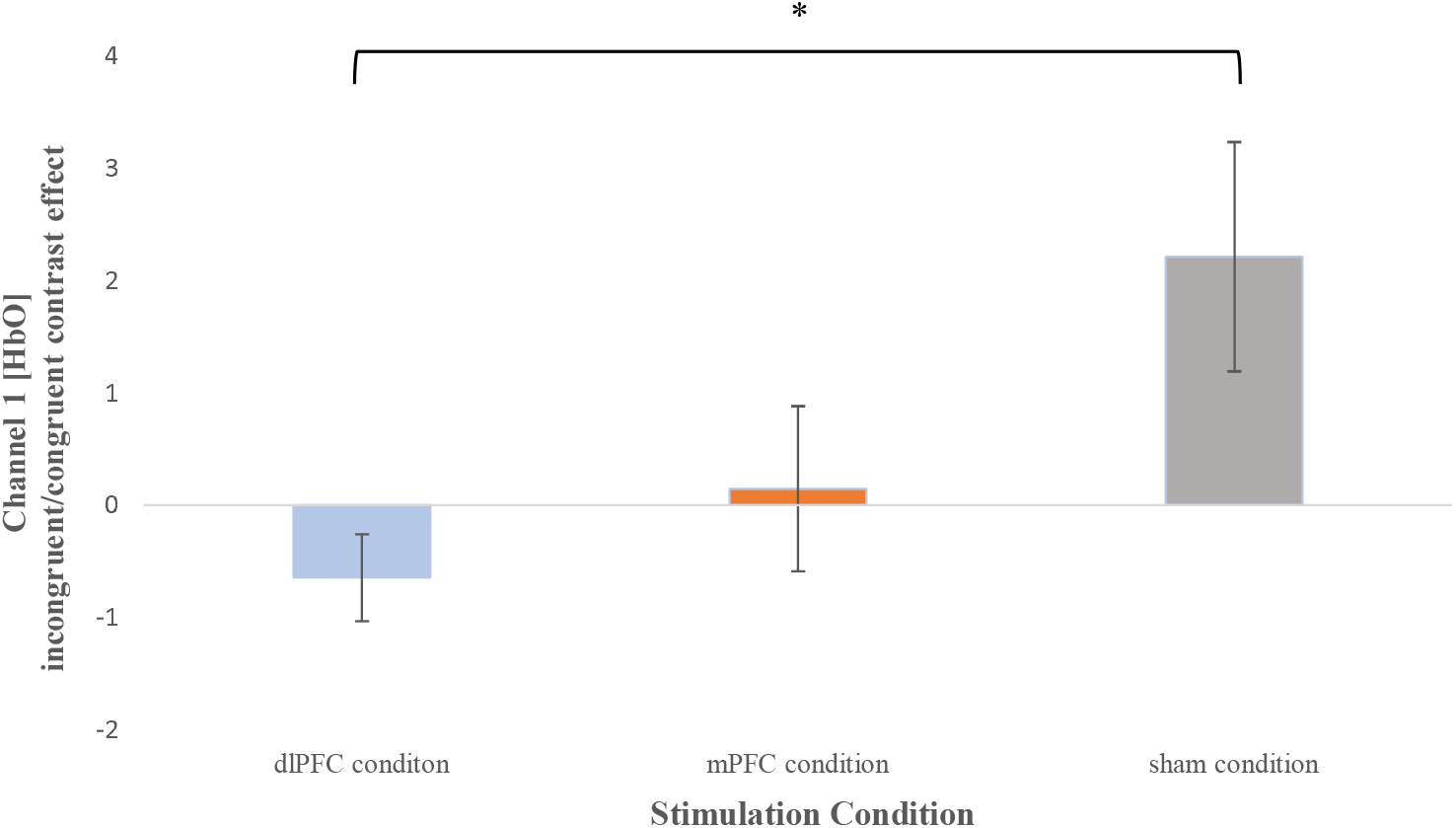
Means (+/-*SE*) for channel 1 oxy-hemoglobin concentration for each stimulation condition on the incongruent/congruent contrast effect; a) dlPFC condition (*M* = −.646, *SE* = .386), b) mPFC condition (*M* = .146, *SE* = .735) and c) sham condition (*M* = 2.21, *SE* = 1.02). *: p < .05.

Planned comparisons for channel 1 indicated that compared to the sham condition, those in the dlPFC condition had significantly lower concentration of oxy hemoglobin (*t*(1,21) = −2.706, *p* = 0.013) on Flanker performance (incongruent – congruent). In contrast, compared to the sham condition, those in mPFC condition had no significant differences in oxy hemoglobin concentrations (*t* (1,18) = 1.576, *p* = 0.133) on incongruent/congruent contrast effect.

In addition, two-way ANOVA’s (stimulation x age category and stimulation x gender) were generated for this channel to determine if the contrast effect was further moderated by age or gender, and to determine if any interaction effects exist.

#### Channel Specific Effects

##### Channel 1

With respect to oxy-hemoglobin concentrations for the Flanker incongruent/congruent contrast effect, the two-way (stimulation x age category) ANOVA revealed a significant main effect of stimulation (*F*(2,26) = 3.699, *p* = .039), but no significant main effect of age category (*F*(1,26) = 3.052, *p* = .092). The interaction between stimulation condition and age category was not significant (*F*(2,26) = .749, *p* = .483).

With respect to oxy-hemoglobin concentrations for the Flanker incongruent/congruent contrast effect, the two-way (stimulation x gender) ANOVA revealed a marginal effect of stimulation (*F*(2,26) = 3.047, *p* = .065), but no significant main effect of gender (*F*(1,26) = .544, *p* = .467). The interaction between stimulation condition and gender was significant (*F*(2,26) = 6.674, *p* = 0.005); the pattern of means suggests differences in the effects of the stimulation on the incongruent/congruent contrast between males and females for channel 1. Variable means for all stimulation conditions by gender for channel 1 have been depicted in Figure 3 below.

**Figure 3.**
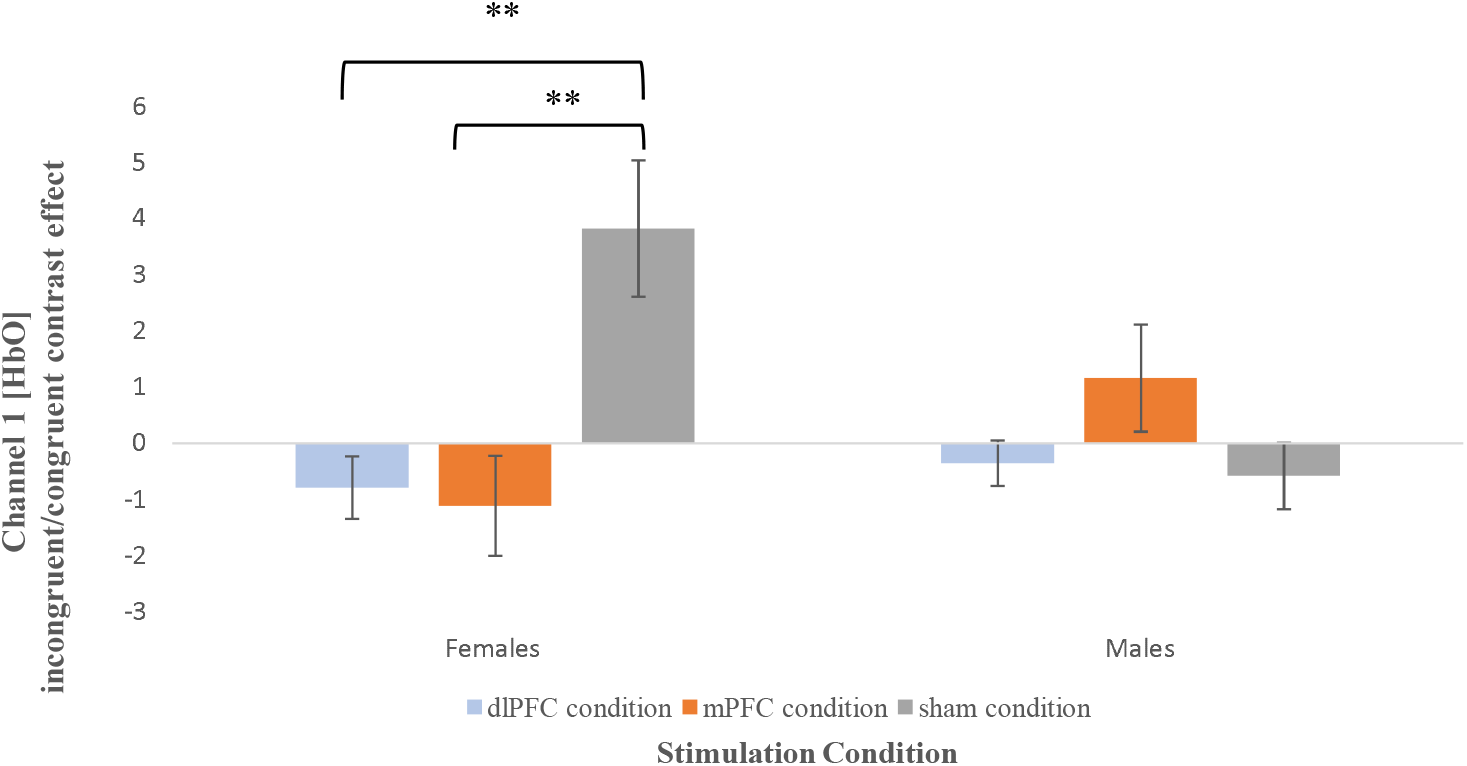
Means (+/-*SE*) for channel 1 oxy-hemoglobin incongruent/congruent contrast effect by gender for each treatment condition; i) Females: a) dlPFC condition (*M* = −.791, *SE* = .553), b) mPFC condition (*M* = −1.114, *SE* = .886) and c) sham condition (*M* = 3.814, *SE* = 1.212); ii) Males: a) dlPFC condition (*M* = −.356, *SE* = .407), b) mPFC condition (*M* = −1.154, *SE* = .950) and c) sham condition (*M* = −.582, *SE* = .593). **: p < .01.

Planned comparisons indicated that compared to the sham condition (*M* = 3.814, *SE* = 1.212), females in the dlPFC (*M* = −.791, *SE* = .553) had a significantly lower incongruent/congruent contrast effect (*t*(1,13) = −3.612, *p* = 0.003). In contrast, compared to the sham condition (*M* = −.582, *SE* = .593), males in the dlPFC condition (*M* = −.356, *SE* = .407) did not show any significant differences on the incongruent/congruent contrast effect (*t*(1,6) = .315, *p* = 0.764).

In addition, compared to the sham condition (*M* = 3.814, *SE* = 1.212), females in the mPFC condition (*M* = −1.114, *SE* = .886) also had a significantly lower incongruent/congruent contrast effect (*t*(1,9) = −2.797, *p* = 0.021). In contrast, compared to the sham condition (*M* = −.582, *SE* = .593), males in the mPFC condition (*M* = −1.154, *SE* = .950) did show have any significant differences on the incongruent/congruent contrast effect (*t*(1,7) = 1.451, *p* = 0.190).

#### Delay Discounting Oxy Hemoglobin Concentration

##### Delay Discounting OxyHb

Following the calculation of the delay discounting oxy hemoglobin concentrations (see Supplementary material), each channel was subject to a one-way ANOVA, to test for any hypothesized group differences (active iTBS vs. sham iTBS). Differences across channels have been illustrated using a heat map and a 3D anatomical overlay (Figure 4), from corresponding *F*-values and *p*-values for each channel.

**Figure 4.**
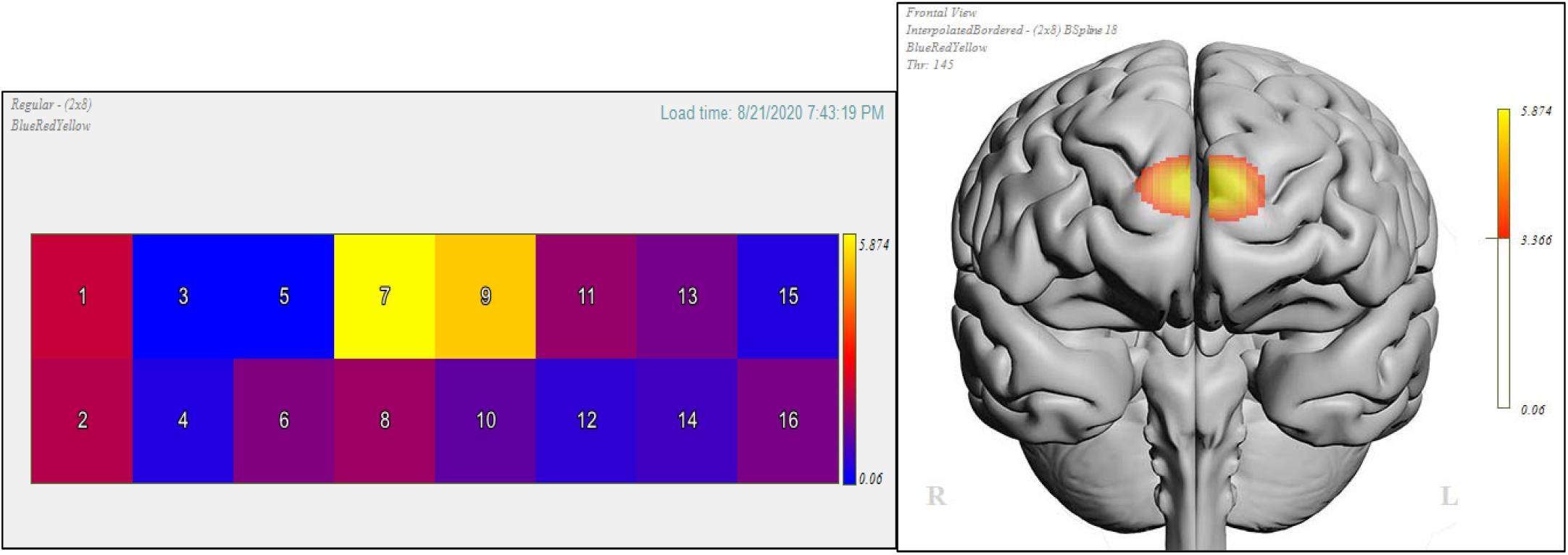
Left: Heat map of channels 1-16 illustrating the difference in oxy-hemoglobin concentration between active iTBS (targeting the bilateral dmPFC) vs. sham iTBS during the delay discounting task. Colour coding was represented by the strength of the p-values; warmer colours represent stronger active vs. sham contrast; darker shades of blue represent weak to no differences. Right: 3D anatomical overlay of the significant contrast difference between active vs. sham.

Analysis revealed channel 7 (*F*(2,34) = 5.874, *p* = .006) and channel 9 (*F*(2,33) = 5.289, *p* = .010) to have significant differences in oxy-hemoglobin concentrations between stimulation conditions (active iTBS vs. sham iTBS). Variable means for both channels by stimulation condition have been depicted in their respective figures below (Figures 5 and 6).

**Figure 5.**
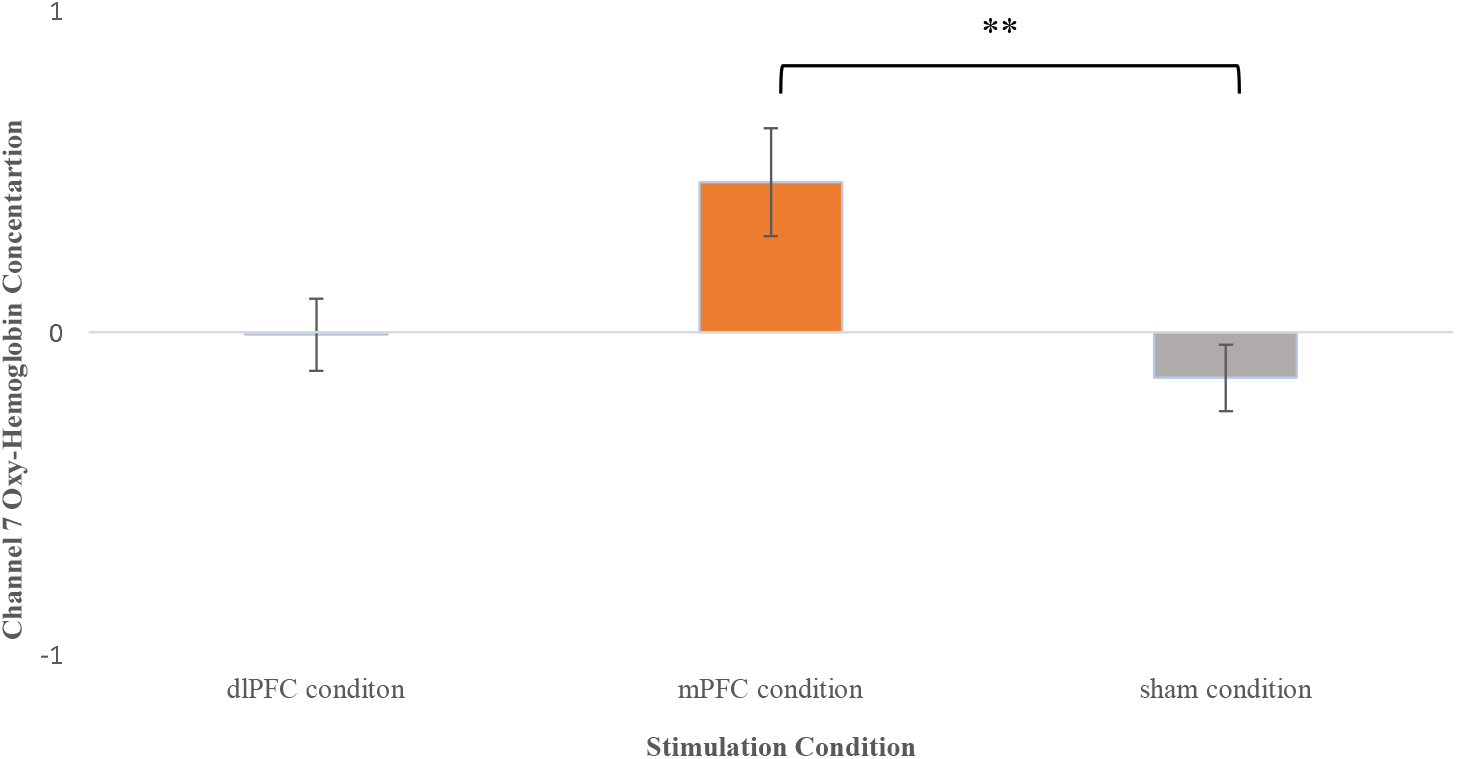
Means (+/-*SE*) for channel 7 oxy-hemoglobin concentration for each treatment condition averaged across the three delay discounting tasks; a) dlPFC condition (*M* = −.00787, *SE* = .112), b) mPFC condition (*M* = .466, *SE* = .167) and c) sham condition (*M* = −.142, *SE* = .103). **: p < .01.

**Figure 6.**
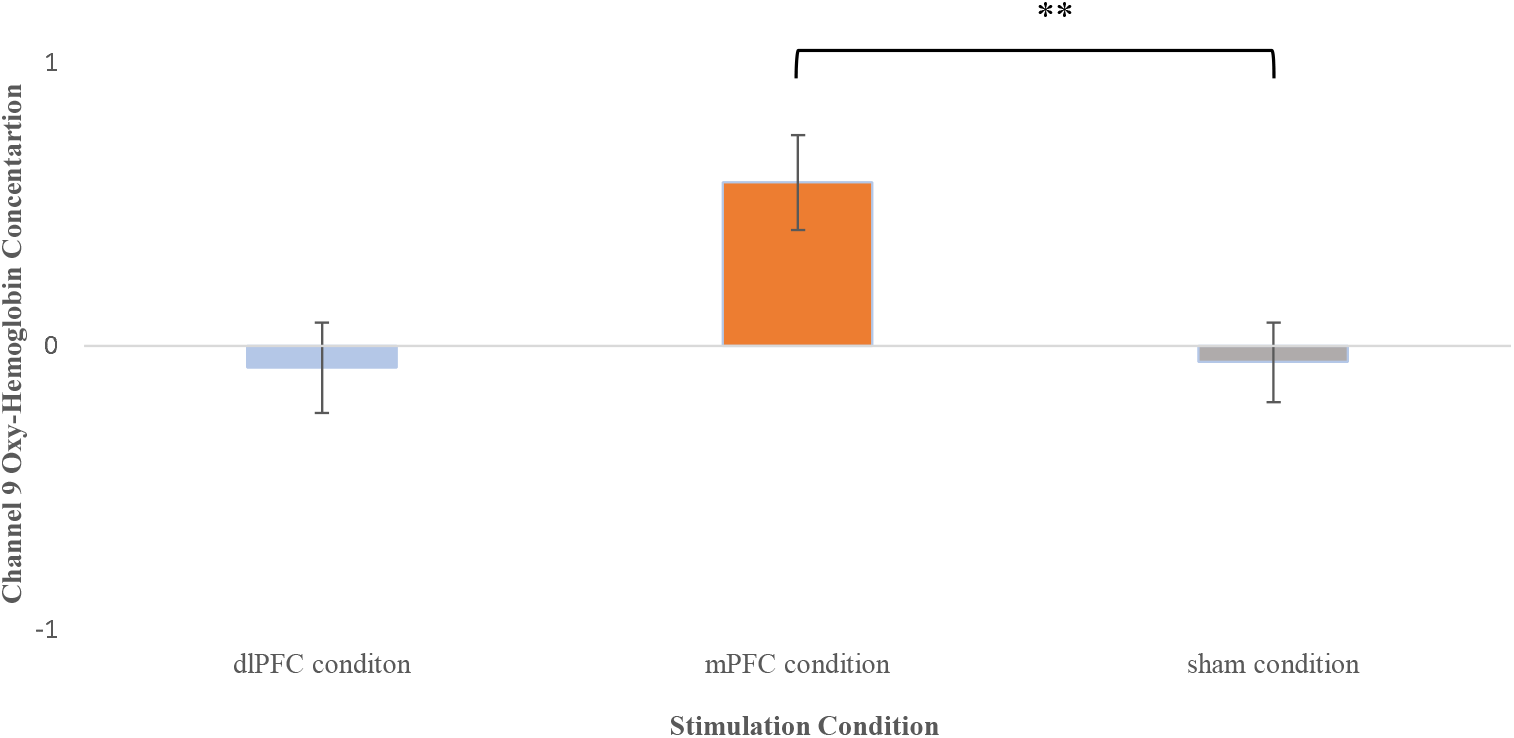
Means (+/-*SE*) for channel 9 oxy-hemoglobin concentration for each treatment condition averaged across the three delay discounting tasks; a) dlPFC condition (*M* = −.0750, *SE* = .160), b) mPFC condition (*M* = .579, *SE* = .167) and c) sham condition (*M* = −.0564, *SE* = .141). **: p < .01.

Planned comparisons for channel 7 indicated that compared to the sham condition, those in the mPFC condition had significantly higher concentrations of oxy hemoglobin (*t*(1,21) = 3.154, *p* = 0.005) on delay discounting task (Figure 5). In contrast, compared to the sham condition, those in dlPFC condition had no significant differences in oxy hemoglobin concentrations (*t*(1,24) = .872, *p* = 0.392) on the delay discounting task.

In addition, a similar pattern was observed for channel 9; compared to the sham condition, those in the mPFC condition had significantly higher concentrations of oxy hemoglobin (*t*(1,20) = 2.902, *p* = 0.009) during delay discounting task (Figure 6). In contrast, compared to the sham condition, those in dlPFC condition had no significant differences in oxy hemoglobin concentrations (*t*(1,23) = −.085, *p* = 0.933) on the delay discounting task.

#### Interaction Analyses

Two-way ANOVA’s (stimulation x age category and stimulation x gender) were generated for both channels to determine if delay discounting oxy hemoglobin concentrations were further moderated by age category or gender. Results revealed no significant interaction effect between stimulation x age category or stimulation x gender for both channels (see Supplementary material for analysis).

### Food Consumption and Task Performance

#### Food Consumption

With respect to food consumption, the two-way (stimulation x age category) ANOVA revealed no significant main effect of stimulation (*F*(2,37) = 0.655, *p* = 0.526) or age category (*F*(1,37) = 3.068, *p* = 0.088). The interaction between stimulation condition and age category was also not significant (*F*(2,37) = 1.231, *p* = 0.304).

With respect food consumption, a two-way (stimulation x gender) ANOVA was conducted to examine the effect of treatment condition and gender on food consumption. The analysis revealed no significant main effect of stimulation (*F*(2,37) = 1.191, *p* = 0.315), but a significant main effect of gender *F*(1,37) = 38.007, *p* < .001) on food consumption. The pattern of means suggests that across study conditions males (*M* = 119.174, *SE* = 8.951) consumed nearly twice as much food as females (*M* = 67.261, *SE* = 4.265). In addition, a statistically significant interaction was found between gender and stimulation group on food consumption (*p* = 0.040, *F*(2,37) = 3.110), suggesting that the effect of stimulation on food consumption was significantly different for males and females. Variable means for all stimulation groups by gender are depicted in Figure 7.

**Figure 7.**
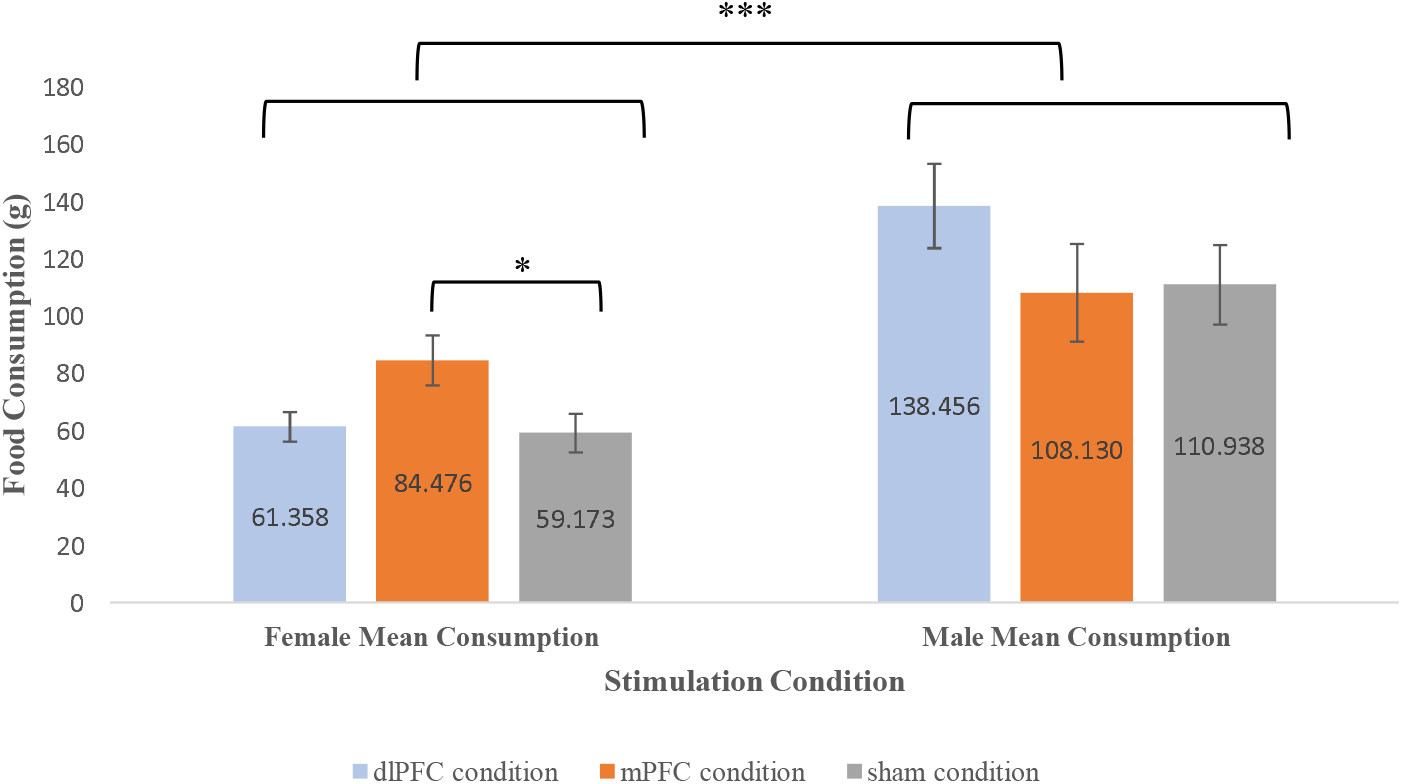
Mean (+/-*SE*) for food consumption (g) by gender for each treatment condition; i) Females: a) dlPFC condition (*M* = 61.358, *SE* = 5.143), b) mPFC condition (*M* = 84.476, *SE* = 8.709) and c) sham condition (*M* = 59.173, *SE* = 6.720); ii) Males: a) dlPFC condition (*M* = 138.456, *SE* = 14.766), b) mPFC condition (*M* = 108.130, *SE* = 17.021) and c) sham condition (*M* = 110.938, *SE* = 13.898). *: p < .05. ***: p < .001.

Planned comparisons indicated that, compared to the sham condition (*M* = 59.173, *SE* = 6.720), females in the mPFC condition (*M* = 84.476, *SE* = 8.709) consumed significantly more food (*t*(1,15) = 2.329, *p* = 0.034). In contrast, compared to the sham condition (*M* = 110.130, *SE* = 13.898), males in the mPFC condition (*M* = 108.130, *SE* = 13.898) did not significantly consume more food (*t*(1,8) = −.128, *p* = 0.901). There were no significant differences in food consumption between those in dlPFC condition and sham condition for both males (*t*(1,8) = 1.357, *p* = 0.212) and females (*t*(1,18) = .263, *p* = 0.796).

Additional analyses were conducted for each food type separately: potato chips (salty) and chocolate (sweet). Two-way ANOVA’s were performed to determine if there were any main effects or interactions between our categorical variables of interest: stimulation, age category and gender (see Supplementary material). Analysis revealed that chocolate consumption was driving the stimulation-induced gender effect.

#### Cognitive Task Performance

Two-way ANOVA’s (stimulation x age category and stimulation x gender) were generated to examine main effects of stimulation group (active dlPFC vs. active mPFC vs. sham stimulation) on Flanker inference score and log transformed delay discounting (*k* values), as well as the interaction effects on each of variable. Results revealed no significant main stimulation effect nor interaction effect between stimulation x age category or stimulation x gender on both cognitive tasks (see Supplementary material).

## Discussion

The purpose of the current study was to examine the effects of excitatory brain stimulation (iTBS) on calorie-dense food consumption, and to examine potential demographic moderators of any such effects. Two prefrontal stimulation targets—the dlPFC and the mPFC— were of specific interest because of their differential roles in executive control and evaluative processing, respectively. Findings revealed reliable effects of iTBS on region-specific increases in neural activity: excitatory stimulation targeting the left dlPFC resulted in a neural efficiency effect in the lateral PFC in response to an interference task, whereas excitatory stimulation targeting the medial PFC resulted in greater neuronal recruitment in response to an evaluative processing task. These effects were theoretically meaningful, and confirm the presence of a stimulation effect on the underlying neural substrates being targeted; the latter is an important finding, as relatively little validation of iTBS effects on the cortex have been undertaken in single stimulation-session format, and no prior studies comparing lateral and medial PFC stimulation using iTBS within the same study exist.

With respect to eating outcomes, findings were mixed. A significant effect of iTBS on eating was evident for females in the dmPFC condition, such that female participants consumed significantly more calorie dense snack foods following active iTBS than following sham stimulation. This reliable, iTBS-induced increase in consumption suggested that enhancement of evaluative processing may have stimulated appetite; such a finding could reflect an amplification of context-induced bias of evaluative processing in the direction of indulgent eating because the presence of generally appetitive foods and a permissive eating environment (e.g., encouragement “to eat as much as you like” to make ratings). An alternative explanation might be that excitatory stimulation of the dmPFC could have generated more spontaneous mind wandering about appetitive dimensions of the food via activation of the default mode network (Horn *et al*., 2014), which would be impelling of consumption particularly under conditions of fasting (as was the case for our participants). Granular analysis by food sub-type revealed that these eating effects were mostly driven by the consumption of sweet snack foods rather than salty snack foods, which may be consistent with this hypothesis.

Sex differences may have emerged due to differences in social cognition and cue sensitivity. It is well documented that females are more prone to higher levels of everyday dietary restraint than men (Rolls *et al*., 1991; Wardle *et al*., 2004; Darcy *et al*., 2013). For this reason, females may have experienced relatively more potentiated evaluative processing in relation to tempting food cues in the eating situation presented. Additionally, fMRI studies have demonstrated higher reactivity in craving and taste-related brain regions to appetitive food cues in woman compared to men (Uher *et al*., 2006; Frank *et al*., 2010; Atalayer *et al*., 2014). Lastly, the eating effect driven by chocolate consumption in females can be explained by previous literature that has identified woman reporting stronger cravings for cues to sweet foods than savory foods, when compared to men (Zellner *et al*., 1999; Hallam *et al*., 2016).

In terms of contribution to the existing research literature, this is the only experimental study to our knowledge that has employed iTBS targeting the dmPFC to assess impact on eating behavior in the laboratory setting. One prior case study (Downar *et al*., 2012) reported the use of high frequency excitatory rTMS targeting the dmPFC increased the ability to control bingeing in bulimia. This perhaps suggests differences in excitatory stimulation effects on the dmPFC between healthy and vulnerable individuals, which could be due to differences in frontal-striatal activity (Downar *et al*., 2012). It should be noted also that the study in question employed multisession stimulation and did not specifically use iTBS. As such, there is ample justification for future research to further explore the effects of neuromodulation of the mPFC and its impact on eating.

The fNIRS data presented in this study highlights the value added by combining neuroimaging techniques with neuromodulation when examining the PFC function in eating research. The findings presented are in agreement with previous studies that have shown neural recruitment of the lateral areas of the PFC to be important for inhibitory/anticipatory mentally effortful tasks (Curtin, Ayaz, *et al*., 2019; Vassena *et al*., 2019), whereas activation of medial PFC has been observed during tasks involving relative weighing of values attached to temporally proximal versus distal outcomes (Marco-Pallarés *et al*., 2010; Mitchell *et al*., 2011; Wang *et al*., 2014). Key strengths of this study include the integration of neuroimaging methods with neuromodulation, the use of blinding combined with sham coil for the sham condition, and the targeting of multiple PFC subregions within the same study. Likewise, the use of a behavioral test of actual food consumption is an important improvement over studies that employ only selfreported measurement of food cravings. Finally, the mPFC is rarely explored as a stimulation target in eating research, and the current study helps to add to this new area of inquiry. Limitations of this study include the relatively modest sample size (though typical of neuromodulation studies), which may have decreased the statistical power for the behavioural measurements. Other limitations of this study include the lack of double blinding and uncertain generalizability; however, our sample did employ a considerably larger age range than most eating studies involving the use of rTMS and/or fNIRS.

In conclusion, the findings from this study suggest that there is a considerable amount of complexity in iTBS effects on eating, and that effects might be modified by sub-region and gender. Findings confirmed expected effects of medial and lateral PFC stimulation on neuronal substrates, but identified a pattern of amplified calorie dense food consumption among those in the mPFC condition compared to sham. This effect was moderated by participant sex, but was relatively invariant across the age span. Future studies would benefit from exploring the range of mPFC stimulation effects that could be produced in inhibitory or facilitating eating environments, and using different variants of TBS protocols (e.g., cTBS). Such studies focussing on the dlPFC have produced interesting findings (Safati and Hall, 2019), and could provide a template for further research involving the mPFC.

## Additional information

The authors IF, AS, NS and PH declare no competing interest.

fNIR Devices, LLC manufactures the optical brain imaging instrument and licensed IP and know-how from Drexel University. HA was involved in the technology development and thus offered a minor share in the startup firm fNIR Devices, LLC. The authors declare that the research was conducted in the absence of any commercial or financial relationships that could be construed as a potential conflict of interest.

All authors have approved this manuscript.

